# On the Ontological Foundations of Cellular Development

**DOI:** 10.1101/2020.05.30.124875

**Authors:** Patryk Burek, Nico Scherf, Heinrich Herre

**Affiliations:** Institute of Computer Science, Faculty of Mathematics, Physics and Computer Science, Marii Curie-Sklodowskiej University, Lublin, Poland; Institute for Medical Informatics and Biometry, Carl Gustav Carus Faculty of Medicine, School of Medicine, TU Dresden, Dresden, Germany; Max Planck Institute for Human Cognitive and Brain Sciences, Leipzig, Germany; Institute for Medical Informatics, Statistics and Epidemiology, University of Leipzig, Leipzig, Germany

**Keywords:** Knowledge management, Ontology of biological reality, Theories of Developmental Biology, Microscopy, Time-lapse imaging, Cell tracking

## Abstract

Time-lapse microscopy is a principal tool to unravel the mystery of how cells form and maintain organisms. The complexity of the domain of cellular dynamics demands a conceptual architecture as a solid theoretical foundation that supports the integration of knowledge obtained across experiments and theories. In this work, we outline the ontological foundation of cellular genealogies, a key concept for describing and representing of cellular development. We build the conceptual framework following the onto-axiomatic method: We first analyse the domain within the context of a top-level ontology (GFO). The resulting domain-specification provides the basis for a conceptualisation where we introduce concepts and relations. From these conceptualisations, we then construct model-structures adhering to the principles of model-theory. We finally elaborate axioms based on these model-structures. The developed framework provides the fundamental concepts underlying a Cell Tracking Ontology (CTO) that supports extraction and integration of biological knowledge from systems-level experiments across different types of observations at the single-cell level.

## 1. Introduction

Cellular dynamics unfolding in space and time organise and shape multicellular life as it develops from a single fertilised egg into a complex organism. After development, cellular processes maintain the organism during its lifetime (tissue homeostasis and regeneration). To fully understand how cells build and maintain structures, we have to be able to observe cellular dynamics and cellular states from experiments (1). One milestone was the reconstruction of the embryonic lineage tree of the nematode Caenorhabditis elegans using microscopy (2). From these roots, modern fluorescence microscopy has turned into a powerful tool to resolve the dynamics of thousands of cells together with readouts of cellular states by fluorescent labels (3). Light-sheet microscopy in particular (4) has enabled us to collect four-dimensional (4D) movies of a range of developing embryos from the fruit fly *Drosophila melanogaster* (5) to mammalian model organisms such as the mouse (6). Complementary to recording cellular dynamics by microscopy, new genetic methods deliver single-cell atlases of gene expression in developing embryos. Those measurements yield detailed information on the genetic state of single cells across development (7) although the resolution in space and time is very coarse. Thus, a critical question in computational biology is how to integrate data from these different experimental modalities (e.g. connect time-lapse imaging with single-cell sequencing) and across experiments (e.g. imaging of the same specimen in several labs) (1). How can we extract knowledge from such data collections? To this end, we also need to develop and refine our concepts and theories to make sense of the intricate patterns we can observe (8,9). As state-of-the-art microscopy becomes widely available to the biology community (10), we need to establish structured and general schemes (11) concurrently to annotate and share the tracking results. We should base these annotations on a solid theoretical foundation: As pointed out in (12) we should regard the underlying terminology and formal concepts themselves as theories about the biological world. Here, we develop the conceptual architecture that supports integration and interoperability in the field of cell tracking experiments. We discuss the concept of *Cellular Genealogy* (13) (or Cell Lineage) as a fundamental notion for the development of the *Cell Tracking Ontology* (14) - an ontology designed for the integration of data obtained from cell tracking experiments.

## 2. The Ontology of Cellular Genealogies

Firstly, we define the notion of cellular genealogy and introduce essential subtypes.

### 2.1. Cell-Collective Genealogies

We consider an individual cell as a material object; hence it has a lifetime, and since cells may divide and eventually die, the number of cells within a region under consideration (e.g. a developing organism) changes through time. Let us consider a time-segment (time-interval) I, such that during I no cell-division and no cell death occurs. Then, the cells existing during I form a collective Coll(I) that can be considered as a continuant through I (15)^2^. During times when the number of cells changes, new cells may occur, and cells may disappear (i.e. die). Let us consider the life of an organism Org from fertilisation to death. Org starts as a single cell, the zygote, develops into a multicellular structure which exists for some time in a dynamic equilibrium (e.g. cells get replenished). After that time, Org dies, i.e. the structures dissolve. We divide the lifetime of Org, LifeT(Org), into a sequence of non-overlapping time-intervals I(1), … I(n) such that the following conditions are satisfied:

1. The intervals I(m) have a first point (they are left-closed), but no last point (right open). More precisely, they have the form [a(m), a(m + 1)) specifying the set {c : a(m) ≤ c < a(m + 1)}, where 0 ≤ m ≤ n. Further, LifeT(Org) = ⋃{I(m) : 0 ≤ m ≤ n}.
2. Let Coll(I(k)) be the set of cells existing during I(k), then no cell death or division occurs during the interval I(k), k ≺ n. Further, we assume that Coll(I(k)) ≠ Coll(I(k + 1)).

These conditions imply further properties: From Coll(I(k)) to Coll(I(k + 1)) the number of existing cells changes. We consider two types: cell division and cell death. If a division of a cell *c* ∈ I(k) occurs, then this process ends up with two daughter cells starting their existence at the left boundary of the interval I(k + 1). Analogously, if a cell undergoes cell death during I(k), then this ends at the left-boundary of I(k + 1). The final definition of CollGen(c(0)) then must specify which cells from Coll(*k*) are related to which cells in Coll(k + 1). To this end, we introduce two relations: div(*x, y, z*): a cell *x* of Coll(*k*) undergoes a cell division during *I*(*k*) resulting in two daughter cells y and z starting their existence at the left-boundary of *I*(*k*+1), and the relation id(*x, y*) stating that *x* belongs to Coll(*k*) and *y* belongs to Coll(*k* + 1) and both cells are identical. We further say that a cell *x* in Coll(*k*) has a successor cell *y* in Coll(k + 1), if *y* is a daughter cell of x or if *y* is identical with *x*, denoted by succ(x, y). The cell collective genealogy CollGen(c(0)), is then specified by the following system CollGen(c(0)) = ({Coll(*k*) | 0 ≤ k ≤ n}, div(*x,y,z*), id(*x,y*)), see Fig. 1a for an example.

**Figure 1.**
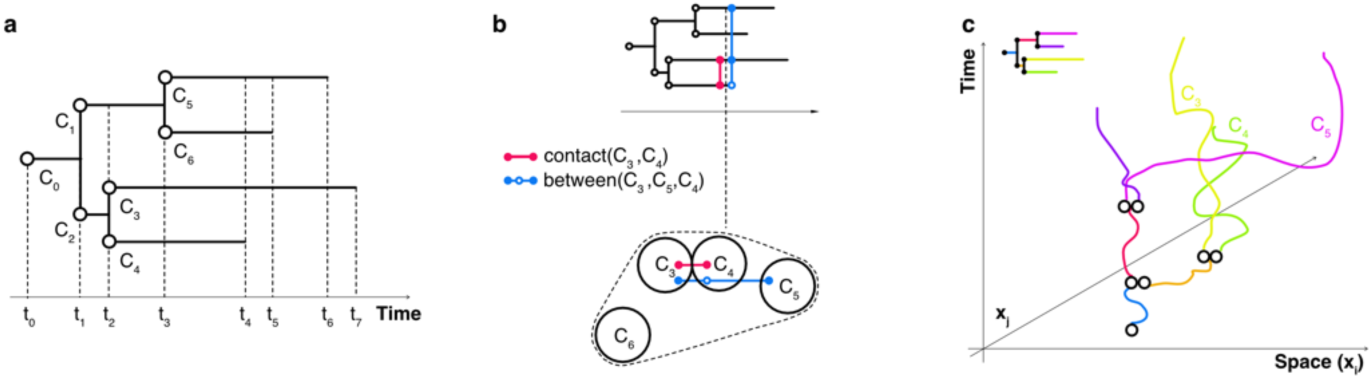
Different conceptual granularities of cellular genealogies: (a) Cell-Collective-Genealogies indicating cell division and cell death as well as the invariance intervals in time. (b) Cell-Situations-Genealogies capture spatial relations between cells in a given context (dashed outline shows convex hull). (c) Cell-Process-Genealogies describe cells as full spatio-temporal processes and their interactions.

We assume that for any organism there exists a uniquely determined cell-collective genealogy. Subsequently, we sketch a proof of this claim. Let t(0) be the starting time-point of the organism (say the zygote). A set of cells, Cells(t), is associated with any time point of the organism’s life, i.e. the set of cells existing at that time-point. Now let t(1) be the latest time-point (> t(0)) such that during the time-interval [t(0), t(1)) no cells are born or die, and at time point t(1) either a cell division or a cell death occurs. t(1) is then the starting point of the next time-interval. The birth or death of a cell marks the end-point of a process, and these processes happen during [t(0), t(1)), and ending in time-point t(1) (death or birth). Assume we have carried out this construction up to time-point t(k). Then we repeat this procedure from t(k) onward, and there is a greatest time-point t(k + 1), greater than t(k), such that during [t(k), t(k + 1)) no cells are born or die, and at t(k + 1) cell birth or death is happening. After a finite number of such steps t(0), …, t(n) we yields a sequence of time intervals, [t(0), t(1)), [t(1), t(2)), …, [t(k), t(k + 1)), …, [t(n - 1), t(n)]^3^ which satisfies the desired conditions^4^.

We conceptually divide the whole life-cycle of an organism (the ontogeny) in various phases (e.g. embryogenesis, growth, ageing) as it develops from a single zygote into its adult form, ages and finally dies (16). As outlined, for every cell collective *x* there is a uniquely determined time-interval I such that no changes occur during *I*. We call this time-interval the *invariance interval* of the cell-collective; it has a left-boundary and no right-boundary. The lifetime of any cell in the collective includes the invariance interval as a temporal part. There is a successor relation between the cell-collectives. The cell-collective *y* is a successor of the collective *x* if the right boundary of the invariance interval of *x* coincides with the left-boundary of the invariance interval of *y*. Further, we group every cell collective with invariance interval I into subsets of Coll(I). During development, certain groups of cells may correspond to the development of specific structures, e.g. organs such as the heart, brain, digestive tract etc.

Understanding the structure of those sub-genealogies is an important research topic in developmental biology. The full sub-genealogy of a particular cell c, being a member of a cell-collective, contains all cells that are in the transitive closure of the successors of this cell. Any cell c generates its cell collective genealogy, denoted by CollGen(c) which is uniquely determined. *Cell lineage* indicates the development history of a tissue, organ or organism from earlier stages such as the fertilised egg.

### 2.2. Cell-Situation-Genealogies (SitGen)

A cell situation genealogy is an extension of a cell collection genealogy: We start with the system CollGen(c(0)) and extend any collective Coll(k) of cells into an object-situation Sit(k). Sit(k) contains precisely the cells of Coll(k) as objects and is embedded into an object-situation with the time-frame *I*(*k*) and a specified space-frame which contains at least the spatial convex closure of the objects in Coll(*k*), see example in Fig. 1b). We are free to add relations between the cells or predicates of the cells in Coll(k). The set of these relations and predicates is inherently open-ended and defines the type of the corresponding situation. A signature Σ determines the situation type. Σ contains the admitted relational and predicate symbols. In the simplest case, this signature Σ is fixed for all situations of the genealogy. It may be necessary to introduce new relations and predicates during development. We consider a cell collective the simplest cell situation as its signature contains only the equality symbol =.

### 2.3. Cell-Process-Genealogies

There are properties of cells, such as velocity or morphodynamics (changes in cell shape) along a trajectory that cannot be attributed to cells as objects and can only be captured by introducing cell processes (cf. (17)). The simplest process genealogy is defined by the transformation of the CollGen into a branching process, denoted ProcCollGen(c(0)). This process is determined by using the integration axiom of GFO and by transforming any cell of Coll(*k*) into a corresponding process. We can then define Proc(*k*) as the integration of all processes Proc(*c*) for any *c* in Coll(*k*). The processes in Proc(*k*) are usually not isolated threads because there can be meaningful interactions between them (e.g. cells exchanging signals by direct contact or diffusible signalling molecules). Hence, there are various versions of potential process genealogies. Analogously, we may transform any cell-situation Sit(*k*) into a process situation, called situoid. The investigation and classification of possible process genealogies is a research field of its own.

We give a simple example to illustrate these ideas in Fig. 1 visualising a part of a cell-collective genealogy. The invariance intervals are: [*t*_*0*_, *t*_*1*_), [*t*_*1*_, *t*_*2*_), [*t*_*2*_, *t*_*3*_), [*t*_*3*_, *t*_*4*_), [*t*_*4*_, *t*_*5*_), [*t*_*5*_, *t*_*6*_), [*t*_*6*_, *t*_*7*_)

The cell collectives, associated with the invariance intervals, are then:

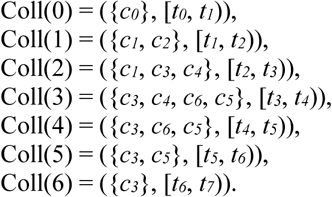

We can extend any of these cell collectives with relations. Here we consider an extension of the cell collective Coll(3). The collective identifies only the different cells contained in it, though it does not specify anything about the relations between those cells. As an example, we introduce the following relations: contact(*x,y*) := the cells *x* and *y* are in contact and between(*x, y, z*) := *z* is between *x* and *y*. The time-frame of Sit(3) coincides with the invariance interval of Coll(3), whereas the space-frame of the situation must be specified additionally (e.g. indicated by dashed region in Fig.1b). Since the cells typically move during the invariance interval, they may change their positions relative to each other (see Fig.1c). The snapshots of such situations are presentic entities, and a situation can be accessed only through its snapshots, which are called *presentic situations*.

A presentic situation cannot adequately describe processes. Let us consider a cell c, as an object having a lifetime and thus, persisting through time: It is the same cell at every time-point of its lifetime. A cell c may move through space, and this movement (trajectory) is a process (Fig. 1c). The presentic locations of c in time is an external attributive of c. The trajectory of a cell cannot be an attributive of c, because the path itself has no presentic nature. If we consider a snapshot of the trace, its form disappears. To model these aspects, we replace any cell in the situation by its corresponding process to get a processual situation.

## 3. Formal Axiomatisation of Cellular Genealogies

In this section, we present a selection of fundamental axioms about cell-collective genealogies. We develop these axioms using the onto-axiomatic method (18) integrating Hilbert’s approach in (19) with a top-level ontology as a general analytical framework. Here we use some axioms from GFO (20) and adapt them to the domain of cell development, and more generally, to the field of developmental biology. We introduce axioms at two levels, the level of GFO-axioms, which are essential to understand the domain-specific notions, and the level of cell biology. Furthermore, we introduce the idea of trans-level axioms connecting concepts across different levels of abstraction.

### 3.1. GFO-Level

#### 3.1.1. Objects and Processes

A material object is a spatio-temporal individual occupying space, persisting through time that is wholly present at every time-point of its lifetime. For every material object *Obj* and a time point *t* of the object’s lifetime, there is a snapshot P of Ob*j* at that time-point *t*, which we express by the expression snapshot (Obj, P, *t*). These snapshots differ from time-point to time-point, though there is something in common, a similarity, between them, which is captured by a universal Univ(*Obj*). Furthermore, any instance of Univ(Obj) stands for Univ(*Obj*). If we consider, for example, an individual cat *C*, and if we take into account Univ(*C*), then any instance of Univ(*C*) stands for the whole universal Univ(*C*). Also, we may imagine a prototypical cat representing this universal. The universal and a corresponding prototype express the phenomenon of persistence, whereas the instances themselves may change their properties through time.

In contrast to an object, an individual process P evolves through time and can never be wholly present at a time-point. The restrictions of P to time-points of P’s temporal extension are called process-boundaries. A process boundary of a process *P* is an entity of presentic nature. However, it can never represent the entire process *P*. Processes and objects thus exhibit a fundamental duality in spatio-temporal reality. Material objects and processes are connected in a particular way, which is expressed by the integration axiom of GFO: For every material object Obj there exists a process Proc(Obj) such that the snapshots of Obj coincide with process boundaries of Proc(Obj). Proc(Obj) is a minimal process associated with Obj, because it exhibits at the process boundaries only such properties that are genuine properties of the object. Genuine properties have a presentic nature and are independent of any process. A process boundary of Proc(*Obj*) contains a snapshot of the object *Obj*. We can extend the minimal process Proc(*Obj*) by adding further properties (that depend on the process) to the process boundaries, e.g. the velocity of a moving object.

#### 3.1.2. Situations

An object-situation (simply called *situation* in this work) is composed of objects that are connected by relators. A situation is framed by a temporal interval and a space region. The material objects contained in a situation Sit define the skeleton of Sit. There is a certain freedom to specify the time frame and the space frame of a situation. For every situation, we may consider snapshots called presentic situations (PSit). An object-situation exhibits a presentic situation at any time-point of its time frame. We generally assume that a presentic entity is a dependent entity. It is either a snapshot of an object (or an object situation), or a part of the time-boundary of a process.

#### 3.1.3 A selection of axioms

We select some axioms as examples, (20) presents a complete system. Axioms are formulated based on signatures providing symbols, predicates and relations.

We now fix a signature Σ(0)^5^ = {Chr(*x*), Obj(*x*), Proc(*x*), Pres(*x*), SReg(*x*), TimeExt(*x*), exhib(*x, y, z*), lifetime(*x, y*), occ(*x, y*), procbd(*x, y, z*), tempext(*x, y*), temprestr(*x, y, z*), tp(*x, y*), where: Chr(*x*):= *x* is chronoid; Obj(*x*) := *x* is an object (in the sense of a material object, being a continuant); Proc(*x*) := *x* is process; Pres(*x*) := is a presential; SReg(*x*) := *x* is a space region; TimeExt(*x*) := *x* is temporally extended entity; exhib(*x, y, z*) := the material object *x* exhibits the entity *y* at time-point *t*; lifetime(*x, y*) := *x* is the lifetime of the object *y*; occ(*x, y*) := *x* occupies space region *y*; procbd(*x, y*) := *x* is a process boundary of the process *y*; procbd(*x, t, y*) := *x* is a process and *y* is the process boundary of *x* at time-point *t*; tempext(*x, y*) := *x* is the temporal extension of the process y; temprestr(*x, y, z*) := *x* is the temporal restriction of the process *y* to the time-interval *z*, being a temporal part of the temporal extension of *y*; tp(*t, x*) := *t* is time-point of the interval *x*. We select some axioms.^6^

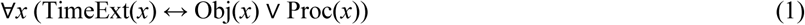

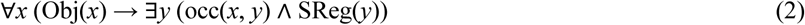

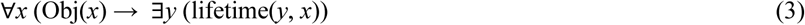

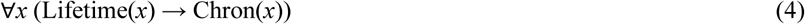

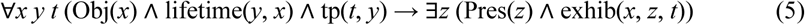

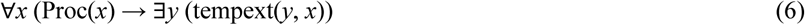

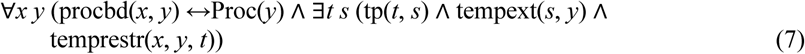

We define the process boundary of a process at time point *t* (being an element of the temporal extension of the process) by the restriction of this process to this time-point.)

We introduce the following integration law^7^. For every material object Obj there exists a process Proc(Obj) such that the snapshots of Obj coincide with the process boundaries of Proc(Obj)). This process exhibits at its boundaries only genuine properties (attributives, i.e. they have a presentic nature and are independent of any process:

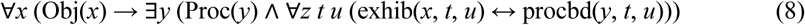

There is a difference between snapshots of objects and process-boundaries: snapshots are taken from objects, never from processes. Presentials have two sources: they can be snapshots of objects (in this case we say that an object *Obj* exhibits a presential at a time-point of its lifetime), or can be contained in boundaries of processes. There are cases when a process boundary is the same as the snapshot of an object participating in this process. In general, the process boundary contains more properties than the process associated with the object. If an object Obj participates in a process *P* then Proc(Obj) is a minimal process layer of P(21). We say that an object Obj participates in a process *P* if any snapshot of Obj is contained within a process boundary of *P*. We introduce the following relation.

partic(*x, y*) := *x* is an object, *y* is a process, and *x* participates in *y*.

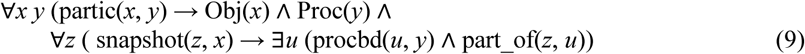

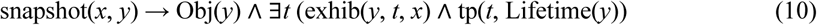

### 3.2. Cell biology level

Cells are considered living entities in contrast to inanimate entities such as stones. However, there is no clear consensus on how to define the boundary between the animate and inanimate. Typical defining properties of life are, among others, metabolism, adaptivity and interaction with the environment, self-organisation, reproduction, heredity, and growth. These conditions define a system which must satisfy at least the following basic properties. It should have a boundary, demarcating the system from the environment, and it should have inner parts. It should further be able to sense and interact with the environment (cf. *Autopoiesis* as an attempt to define living matter using concepts from general systems theory such as self-organisation). In biology, the cell is the simplest system satisfying these assumptions. It is an open problem whether these conditions - though necessary for the definition of life – are also sufficient for determining the essence of the animate. A minority of the biologists believed that an additional life force is needed to achieve a complete picture of the world (22). The self-organised development of a cellular genealogy, starting from a zygote, seems to be an essential feature of the animate. Hence, the ontology of biology should consider the existence of cellular genealogies as one of the basic features demarcating biology from other fields of natural science, as physics or chemistry. Thus, we include relevant concepts of cellular genealogies, such as Cell(*x*), Coll(*x*) cell collective, cell division, cell death, cell situation Sit(*x*) and the corresponding processes in the basic notions of life. The formalization of these notions use the signature Σ(1) = {Cell(*x*), Coll(*x*), CollGen(*x*), Sit(*x*), PSit(*x*), Dead(*x*), id(*x, y*), div(*x, y, z*), invar(*x, y*), member_of(*x, y*), daughter_of(*x, y*), invar(*x, y*)}, where: Cell(*x*) := *x* is a cell; Coll(*x*) := *x* is a cell collective; and CollGen(*x*) := *x* is a cell-collective genealogy.

A cell-collective has members, member_of(*x, y*) := the cell *x* is member of the collective *y*. Its invariance interval determines the lifetime of a collective: inv(*x, y*) := *x* is the invariance interval of the collective *y*. We distinguish two kinds of Time-Entities: Time Points and Time Intervals, where a time point is an element of a time interval. We use two types of time-intervals, those which are closed (they have a first point and a last point), and such which are left-closed and right open (i.e. they have a first point, but no last point). Notable examples are cell-division, cell-death and the various structural and morphological properties of cells.

Subsequently, we present a selection of axioms.^8^

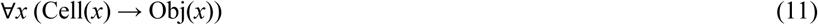

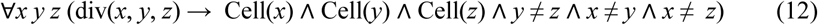

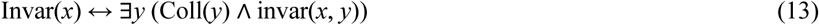

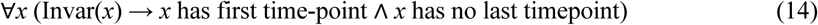

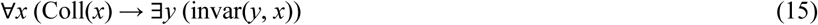

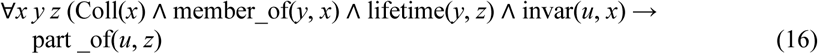

A cell situation, Sit, contains a cell collective forming the skeleton of the situation, and various relations between cells, called the situation’s signature. As an example of a signature consider Σ = (contact(*x, y*), between(*x, y, z*), equidistance(*x, y, u, v*)).

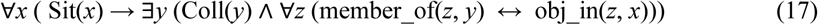

For every situation S there exist a cell-collective C such that the members of C are exactly the objects in S. Here we assume that the situations are spanned by the cells of a collective.

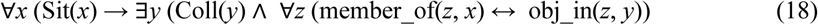

Since the cells of a situation can move during the situation’s time-frame the relations between them may depend on time, e.g. two cells c, d are in contact at time point *t*, and separated at another time-point *t’*. Hence, the relations (e.g. contact(*x,y*)) must be extended by a time-argument such as contact(*x, y, t*). The time-frame of situation *S* is the invariance interval of the collective contained in *S*. coll_succ (*x, y*) denotes a successor relation such that *x* and *y* are cell collectives, and *y* is the successor of *x*.

There exists exactly one cell-collective without a predecessor and exactly one cell collective without a successor. A cell-collective genealogy CollGen is a temporally extended structure consisting of a sequence of invariance intervals and the cell-collectives associated with these intervals: CollGen = (Coll(1), …, Coll(m), inv(1), …, inv(*m*), coll_succ(*x, y*), id(*x, y*), div(*x, y, z*), Dead(*x*)). A sequence of intervals specifies such a cell genealogy Int(CGen) =(inv(1), …, inv(*m*)), by the collectives Coll(*i*), with added successor relation, and (at least) two relations id(*x, y*), and div(*x, y, z*) between the cells of a collective, and the cells of the successor collective. By adding relations to the cell-collectives of a genealogy, we define the notion of a cell-situation genealogy, denoted by SitGen. A situation genealogy is said to be stable if the signature is the same for any situation of the genealogy. A many-sorted model-structure of a cell-collective genealogy can be specified as follows: CollGen = ((L,Inv(1),…,Inv(*n*), <), Cell, Coll(1), …, Coll(*n*), lifetime(*x, y*), succ(*x, y*), id(*x, y*), div(*x, y, z*)).

Here, Cell(*x*) is a predicate, the extension^9^ of which contains all cells occurring during the full temporal extension of the genealogy. The extension of Coll(*i*) are subsets of the extension of the predicate Cell(*x*). (*L*, <) is a dense linear ordering, presenting the set of time-points, and Inv(i) are left-closed and right-open intervals of (*L*, <).

By adding further relations, presented formally by a signature Σ = (r(1), …, r(n)), we get a model-structure for a situation genealogy SitGen: SitGen = (CollGen, int(Σ)). int(Σ) is the interpretation of the relational symbols of Σ in the corresponding cell-collectives Coll(*i*)^10^.

#### 3.2.1. Description of the relations

We introduce the following relations: lt(*x*) := lifetime of the cell *x* and is defined by the following condition lt(*x*) = *y* ⟷ lifetime(*x, y*); daughter(*x, y*) := ∃*z* (div(*x, y, z*); Inv(i)(*x*) := *x* is an element of the i-th invariance interval; Init(*i*)(*x*) := *x* is the initial time-point of the interval Invar(*i*); Init(*x*) := ∨{Init(i)(*x*) | *i* ≤ *n*}; init(*x, y*) := *x* is a cell and *y* is the initial time point of the cell’s lifetime; Coll(*i*)(*x*) := *x* is an element of the *i*-th cell collective. The definition of succ(*x, y*) uses the following formulas: φ(i)(*x, y*) := *x* ∈ Coll(i) ∧ *y* ∈ Coll(i + 1) ∧ (id(*x, y*) ∨ (daughter(*x, y*)); then succ(*x, y*) := ∨{φ(i)(*x, y*) | 0 ≤ i ≤ n -1}.

#### 3.2.2. Selection of axioms

(*L*, <) is a dense linear ordering. Inv(*i*), *i* = 1,…,*n*, are intervals, such that the following conditions are satisfied:

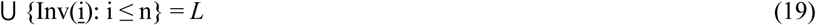

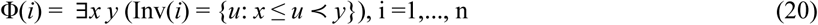

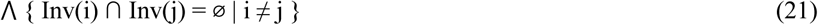

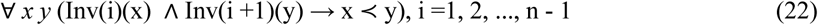

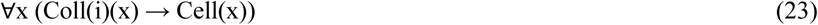

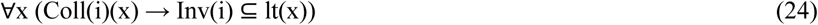

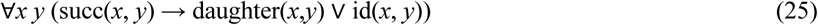

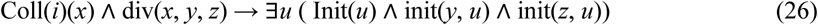

A cell situation genealogy SitGen is based on a cell-collective genealogy CollGen extended by adding relations to any of the cell-collectives.

## 4. The experimental framework and its Formalisation

### 4.1. Basic conditions - Ontology of Frame-Sequences

In this section, we investigate and analyse cell tracking experiments based on the principle of time-lapse microscopy. In reality (*in vivo* as well as *in vitro*) cells are moving and changing continuously in time and space. Hence, the time-points are densely ordered: after a time-point, there is no direct successor. In the considered experiments, discrete snapshots of the continuous dynamics are taken. These snapshots provide incomplete information about an individual situation genealogy of the independent reality. Let a given situation genealogy SitGen be specified by the structure SitGen = ((L, Inv(1), …, Inv(*n*), <), Sit(1),…, Sit(*n*), lt(*x,y*), succ(*x, y*), id(*x, y*), div(*x, y, z*), int(Σ)). The time-points at which the snapshots are taken from a finite subset S ⊆ L of the linear ordering (L, <), hence (S,<) is a finite linear ordering which can be ordered by natural numbers. A snapshot at time point t yields a presentic situation PSit(t), which is called the frame at t, denoted by Fr(*t*). Any experiment Exp of this type results in a finite sequence Seq(SitGen)) = (Fr(t(1)),…, Fr(t(n)) of frames, called components of the sequence. This sequence, related to an experiment Exp, is denoted by Seq(Exp). We say that a time-lapse experiment Exp is adequate for the situation genealogy SitGen if for any situation Sit in SitGen there exists a snapshot of Sit in Seq(Exp).^11^ These sequences are the entities to be investigated. Any of the pictures Fr(k) reflects a snapshot of a situation from SitGen. For the sake of simplicity, we identify the frame Fr(i), being a picture, with the snapshot of the reflected situation. In the following, we fix a sequence Seq(Exp) as a result of a certain experiment.

Every frame is a snapshot of a situation. Hence a frame is a presentic situation (PSit). Further, any presentic situation in FrSeq contains presentic cells, also called presentials. Presentials possess various properties and can relate to other entities. Some properties are inherent to the objects, e.g. the form (based on metrics) or the number of proteins of a certain type. Others are external to the cells, such as the distance between two cells, and the position of the cell in space. Further important relations between two presentic cells are: contact(*x, y*) := the cells I and y are in contact; or relative spatial positions between the cells *x* and *y*, for example, the cell *y* is right to the cell *x, y* is left to *x*, above, below or spatial relations with respect to an (often anatomical) frame of reference (e.g. dorsal, ventral, distal etc…). Also, spatial relations with more than two arguments are possible, e.g. between(*x, y, z*) the cell *y* is localised between the cell *x* and the cell *z*. A further example of a relation with four arguments is equidist(*x, y, z, u*) meaning that the distance between *x* and *y* is the same as that between *z* and *u*.

Although this set of spatial relations may seem quite limited, from a biology point of view, it is in itself already useful to describe a large class of symmetries that are established or broken (24) during development (e.g. mirror symmetry in bilateral animals). Further, from a theoretical perspective, the relations of betweenness and equidistance are even sufficient to establish the whole elementary planar Euclidean geometry (25). We emphasise that in a single frame, only presentic properties can be identified, that are independent of any process. Aspects such as the circularity of a cell path or the morphodynamics of a cell (its particular pattern of shape changes) cannot be detected in any one individual frame and are thus not presentic, but processual properties. The derivation of a processual property of a cell or a cell-collective can only be achieved by an analysis of a sequence of frames. In essence, a sequence of frames can be transformed into a video providing processual properties. Since the famous works of the photographer Eadweard Muybridge to study motion from a sequence of static pictures, the method of changing time scales via slow-motion or time-lapse/fast motion provided many insights in the processual properties of nature, and in particular into the properties of embryogenesis (8,9).

### 4.2 Formal Axiomatisation of frame sequences

#### 4.2.1 Some predicates and informal description of axioms

FSeq(*x*) denotes a frame-sequence *x*, and its components are called frames. Every frame is a snapshot of a situation, denoted by PSit. We introduce a linear ordering between the components of a frame-sequence, hence such a sequence can be presented by the structure FSeq = ({F(1), …, F(*n*)}, <), where F(1) < … < F(*n*). Let Seq be a frame sequence; we say that a component G of Seq is a successor of the component F of Seq, if F < G and there is no component between F and G. We assume that in any frame there occur cells, that these cells are presentials, and any such presentic cell is a snapshot of a uniquely determined cell (with lifetime> 0).

#### 4.2.2 Selection of formal axioms

We first introduce a signature Σ(2) on which the axioms are based: Fr(x) := x is a frame, FSeq(x) := x is a frame sequence, PCell(x):= x is a presentic cell, PSit(x) := x is a presentic situation, comp(x,y) := x is a component of the frame-sequence y, < is the linear ordering between the components of frame sequence, ipart(x,y) := x is an image-part of the frame y.

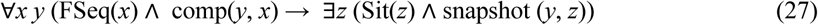

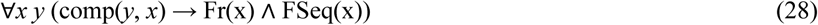

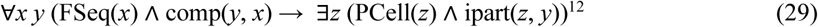

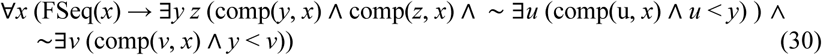

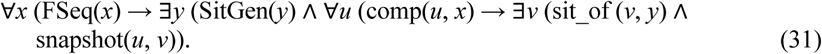

Axiom (31) establishes a link between the experiment and the independent reality of situational genealogies. Such an axiom should be postulated for any type of experiment as each experiment is directed at objects to be studied.

We have established a relation between cellular genealogies and sequences of frames from time-lapse experiments. The final reconstruction of genealogies is then an information artefact that captures relevant knowledge about the real-word genealogies.

## 5. Conclusion and Future Research

In this work, we outlined an ontological foundation of cellular genealogies concerning a fundamental theory and a formal representation of a type of experiments and its results. The full framework will provide three levels of abstraction. This paper addresses the first two levels: the theory and the experiment level. At the *theory level*, we analysed cellular genealogies as independent real-world entities using the onto-axiomatic method. We proposed a partial formal axiomatisation of knowledge assumed to be true for every cellular genealogy. At the *experiment level*, we formally described time-lapse experiments and developed an axiomatic foundation of this domain. Any experimental framework should be considered as a mediator between a theory and the real-world entities to be studied. Experiments provide data about a domain of interest; they play an indispensable role for supporting or disproving a theory, and thus for further development and revisions of theories. Our development of the overarching conceptual framework follows the onto-axiomatic method adhering to the principles of model-theory (23,26) as introduced in biology by (27).

As a conceptual next step, we will extend the genealogy types outlined here, to model the self-organising processes in biology as complex, interacting systems (embryonic development being a prime example). Building on existing work on collective phenomena by (15), we will consider groups of cells and their mutual interactions. We could model groups of interacting cells^13^ as object-situations, e.g. a cell-group can be a specific tissue (or a group of precursor cells). We may further introduce material boundaries and dynamics of these cell-groups to build a cell-group genealogy. A critical problem will be to find appropriate levels of granularity. Here, we will build on ideas from complex systems research.

Finally, we outline directions for future research, as we feel that the presented framework paves the way for new questions and might even open new fields:

### Development of suitable representational levels

We presented theories on a general level in this paper. However, the instance-level is still needed, if we want to study individual cellular genealogies. We are currently investigating various representational levels as continuation of the present paper.

### Extending the ontological foundation of cellular genealogies

To elaborate on the presented framework, we will analyse existing knowledge in developmental biology and successively transform it into formalised axioms based on the onto-axiomatic method.

### Elaborating Genealogy-Theories for particular model species

An ideal first step would be the development of a complete genealogy-theory for the model organism *Caenorhabditis elegans* as much is known about its genetics and development (2,28).

### Extending the outlined theory to other levels of granularity

Our current genealogy-theory refers to the single-cell level as a ‘middle-out’ starting point as already proposed by (29)^14^. We will consider two canonical extensions of granularity levels: We explicitly model the state of cells at the molecular level using (30,31) and we model cell-groups as tissue-level entities.

### Modelling cellular genealogies in disease

To support computational approaches in systems medicine, we should elaborate a specific theory for abnormal genealogical patterns as can be found in certain cancers, such as leukaemia (32) and related diseases.

## Footnotes

2 We stipulate that a cell collective Coll preserves the number and the identity of the cells contained in Coll.

3 With the exception of the last interval all other intervals are left-closed and right-open.

4 We assume that the time-points are represented by real numbers. A complete proof of the conditions uses the continuity of real numbers, in particular the fact that a bounded set of real numbers has a least upper bound.

5 Σ(0) is a minimal signature for the material ontological region according to GFO and must be extended in various directions. We consider biology as belonging to the material stratum of reality.

6 Establishing a fully developed system of axioms is a research topic of its own. Here we present only a selection of particularly important axioms.

7 The integration law is a unique condition distinguishing GFO from other current top level ontologies.

8 A more complete axiomatization of cellular genealogies is work in progress.

9 The extension of a predicate P(*x*) is the set of all entities satisfying this predicate. This notion can be explicated based on a model-structure established according to the methods of logic and model theory (23).

10 If r(*x, y, z*) is a ternary relation symbol in Σ, then an interpretation of *r*, denoted by int(*r*), in Coll(*i*) = {a(1),, a(*n*)} is a subset of Coll(*i*) × Coll(*i*) × Coll(*i*) (e.g. the relation between(*a, b, c*)).

11 This condition implies that the temporal distances between the snapshots are sufficiently small to acquire the relevant information about a cell-division.

12 There is an ambiguity between *part of* a frame and *image-part* of a frame. For sake of simplification we do not distinguish between the image of an entity and the entity itself. We could simply say that a presentic cell is a part of a frame. Though, a frame can possess image parts as artefact to which no real entity corresponds.

13 Such as the parasegments forming during patterning of Drosophila embryos (16).

14 Sidney Brenner is being credited with saying ‘I believe very strongly that the fundamental unit, the correct level of abstraction, is the cell and not the genome’ by (30).

## References

[1] Wallingford JB. The 200-year effort to see the embryo. Science. 2019 Aug 23;365(6455):758–9.

[2] Sulston JE, Schierenberg E, White JG, Thomson JN. The embryonic cell lineage of the nematode Caenorhabditis elegans. Dev Biol. 1983 Nov;100(1):64–119.

[3] Megason SG, Fraser SE. Imaging in systems biology. Cell. 2007 Sep 7;130(5):784–95.

[4] Huisken J, Swoger J, Del Bene F, Wittbrodt J, Stelzer EHK. Optical sectioning deep inside live embryos by selective plane illumination microscopy. Science. 2004 Aug 13;305(5686):1007–9.

[5] Royer LA, Lemon WC, Chhetri RK, Wan Y, Coleman M, Myers EW, et al. Adaptive light-sheet microscopy for long-term, high-resolution imaging in living organisms. Nat Biotechnol [Internet]. 2016 Oct 31;

[6] McDole K, Guignard L, Amat F, Berger A, Malandain G, Royer LA, et al. In Toto Imaging and Reconstruction of Post-Implantation Mouse Development at the Single-Cell Level. Cell [Internet]. 2018 Oct 11

[7] Harland RM. A new view of embryo development and regeneration. Science. 2018 Jun 1;360(6392):967–8.

[8] Wellmann J. Model and movement: studying cell movement in early morphogenesis, 1900 to the present. Hist Philos Life Sci. 2018 Sep 11;40(3):59.

[9] Wellmann J. Die Form des Werdens: eine Kulturgeschichte der Embryologie; 1760-1830. Wallstein; 2010.

[10] Power RM, Huisken J. Putting advanced microscopy in the hands of biologists. Nat Methods [Internet]. 2019 Oct 16;

[11] Gonzalez-Beltran AN, Masuzzo P, Ampe C, Bakker G-J, Besson S, Eibl RH, et al. Community Standards for Open Cell Migration Data [Internet]. bioRxiv. 2019 [cited 2019 Dec 16]. p. 803064.

[12] Leonelli S. The challenges of big data biology. Elife [Internet]. 2019 Apr 5;8.

[13] Glauche I, Lorenz R, Hasenclever D, Roeder I. A novel view on stem cell development: analysing the shape of cellular genealogies. Cell Prolif. 2009;42(2):248–63.

[14] Burek P, Scherf N, Herre H. A pattern-based approach to a cell tracking ontology. Procedia Comput Sci. 2019 Jan 1;159:784–93.

[15] Wood Z, Galton A. A taxonomy of collective phenomena. Appl Ontol. 2009;4(3–4):267–92.

[16] Wolpert L, Tickle C. Principles of Development. OUP Oxford; 2011. 616 p.

[17] Burek P, Scherf N, Herre H. Ontology patterns for the representation of quality changes of cells in time. J Biomed Semantics. 2019 Oct 16;10(1):16.

[18] Baumann R, Loebe F, Herre H. Axiomatic theories of the ontology of time in GFO. Appl Ontol. 2014;9(3– 4):171–215.

[19] Hilbert D. Axiomatisches Denken. In: Hilbert D, editor. Dritter Band: Analysis · Grundlagen der Mathematik · Physik Verschiedenes: Nebst Einer Lebensgeschichte. Berlin, Heidelberg: Springer Berlin Heidelberg; 1935. p. 146–56.

[20] Herre H. General Formal Ontology (GFO): A Foundational Ontology for Conceptual Modelling. In: Poli R, Healy M, Kameas A, editors. Theory and Applications of Ontology: Computer Applications. Dordrecht: Springer Netherlands; 2010. p. 297–345.

[21] Herre H, Heller B, Burek P, Hoehndorf R, Loebe F, Michalek H. General Formal Ontology (GFO): A Foundational Ontology Integrating Objects and Processes. Part I: Basic Principles (Version 1.0). Research Group Ontologies in Medicine (Onto-Med), University of Leipzig; 2006.

[22] Driesch H. Philosophie des organischen; Gifford-vorlesungen gehalten an der Universität Aberdeen in den jahren 1907-1908,. 1921

[23] Hodges W, School of Mathematical Sciences Wilfrid Hodges, Wilfrid H. Model Theory. Cambridge University Press; 1993. 772 p.

[24] Neville AC. Animal asymmetry The Institute of Biology’s Studies in Biology. London, UK: Edward Arnold. 1976;

[25] Tarski A. What is elementary geometry? In: Studies in Logic and the Foundations of Mathematics. Elsevier; 1959. p. 16–29.

[26] Chang CC, Keisler HJ. Model Theory. Vol. 73. Elsevier; 1990.

[27] Woodger JH, Floyd WF. The Axiomatic Method in Biology, By J.H. Woodger. With Appendices by Alfred Tarski and W.F. Floyd. Cambridge University Press; 1937. 174 p.

[28] Brenner S. Nature’s gift to science (Nobel lecture). Chembiochem. 2003 Aug 4;4(8):683–7.

[29] Noble D. The music of life: biology beyond the genome. Oxford: Oxford University Press; 2006.

[30] Bard J, Rhee SY, Ashburner M. An ontology for cell types. Genome Biol. 2005 Jan 14;6(2):R21.

[31] Ashburner M, Ball CA, Blake JA, Botstein D, Butler H, Cherry JM, et al. Gene ontology: tool for the unification of biology. The Gene Ontology Consortium. Nat Genet. 2000 May;25(1):25–9.

[32] Bahr C, von Paleske L, Uslu VV, Remeseiro S, Takayama N, Ng SW, et al. A Myc enhancer cluster regulates normal and leukaemic haematopoietic stem cell hierarchies. Nature. 2018 Jan 25;553(7689):515–20.

